# FarmCPUpp: Efficient Large-Scale GWAS

**DOI:** 10.1101/238832

**Authors:** Aaron Kusmec, Patrick S. Schnable

## Abstract

Genome-wide association studies (GWAS) are computationally demanding analyses that use large sample sizes and dense marker sets to discover associations between quantitative trait variation and genetic variants. FarmCPU is a powerful new method for performing GWAS. However, its performance is hampered by details of its implementation and its reliance on the R programming language. In this paper we present an efficient implementation of FarmCPU, called FarmCPUpp, that retains the R user interface but improves memory management and speed through the use of C++ code and parallel computing.

## Introduction

Genome-wide association studies (GWAS) use statistical tests to identify associations between genetic markers and phenotypes. Next-generation sequencing and improved phenotyping methods have enabled the use of increasingly large datasets in GWAS, posing challenges for the statistical models used and their computational efficiencies.

Two challenges arise from the biology underlying quantitative trait variation. One challenge is false negative marker-trait associations, which are often attributable to small sample sizes and the small effect sizes expected from most QTN. This challenge is met primarily by increasing sample sizes and through experimental designs that increase the precision of phenotypes. The other challenge is false positive marker-trait associations that can arise due to linkage between markers and quantitative trait nucleotides (QTN) induced by population and family structure. This challenge has been addressed through more sophisticated single-marker models that incorporate subpopulation assignments (STRUCTURE) [1], principal components of the marker matrix [2], and kinship matrices [3]. Through this work the general linear model (GLM) and mixed linear model (MLM) [3] became the dominant statistical paradigms in GWAS.

Significant new ideas were introduced by the multi-locus mixed model (MLMM) [4] and SUPER [5] methods. The MLMM addressed the issue posed by linkage disequilibrium between nearby markers by using model selection and multiple regression in conjunction with the MLM. SUPER introduced a new formulation of the kinship matrix based on pseudo-QTNs that are selected from a preliminary GWAS step. However, more complex statistical methods also impacted the computational efficiency of these methods.

The sizes of GWAS datasets also pose a challenge for the computational efficiency of the statistical methods. Various statistical and computational techniques, such as EMMAX [6], P3D [7], and Fast-LMM [8], have steadily improved the computational efficiency—but not the statistical power—of newer software relative to the original implementation of the MLM.

FarmCPU [9] is an innovative method that combines the model selection strategy of MLMM with the restricted kinship matrix of SUPER. Briefly, FarmCPU implements the GLM with optional population structure covariates to scan single markers. The results of this scan are used to select multiple sets of pseudo-QTNs similar to SUPER. The best set is selected based on its log-likelihood estimated from a random effects model. Markers are scanned again using the GLM plus pseudo-QTNs as covariates, and the process is repeated until the set of pseudo-QTNs does not change.

FarmCPU has been reported to be not only a statistically powerful but also a computationally efficient method for GWAS. However, the performance tests reported in the main text of the paper tested at most 60,0 markers with a sample size of 11,000 (66 million data points) (Figure 6 in [9]). Tests on larger datasets were reported in the supplementary materials. Of particular interest to us is the test on 10,000 individuals and 1,000,000 markers (10 billion data points) reported in Figure S26. This figure reports that FarmCPU completed the analysis in ~4 hours on a laptop with 4 GB of RAM. We initially tested FarmCPU with a dataset of 4,890 individuals and 2,452,207 markers (~12 billion data points) and found that this required 9 hours and 265 GB of RAM. This disparity is not due to the use of inferior hardware.

We had a large number of GWAS to perform such that existing software (e.g., MLMM, SUPER, FarmCPU) would have taken prohibitively long. Therefore we investigated the source code of FarmCPU to identify memory- and time-related bottlenecks that could potentially be improved.

## Materials and Methods

### Computational Considerations for GWAS

The sizes of GWAS datasets pose memory management and computational speed challenges for software development. Univariate GWAS requires at minimum a *n* × 1 vector of phenotypes and *n* × *m* marker matrix, where *n* is the number of samples and *m* is the number of markers. A *n* × *q* matrix of covariates, where *q* is the number of covariates, containing potential confounding variables, such as population structure, may also be included. The values of both *n* and *m* have increased rapidly as larger samples are collected to increase statistical power and whole-genome resequencing identifies more variants. Thus, how the marker matrix is represented and accessed in memory is important for all GWAS methods.

While FarmCPU shares the challenge of memory management with other GWAS models, its efficiency concerns differ. Popular early methods, such as the general linear model (GLM) and mixed linear model (MLM), tested each marker once in an analysis. Researchers devised multiple algorithms to improve the efficiency of the MLM, although none of these improved its statistical power. While FarmCPU uses the efficient GLM, it also uses an iterative, model selection approach to increase statistical power and decrease false positives. This requires testing each marker multiple times, adding to the computational burden of the analysis.

Subsequent sections discuss three ways in which FarmCPUpp improves on the original implementation of FarmCPU through the use of better memory management techniques, compiled code, and parallel computing.

### Memory management

R’s design makes working with even modest numbers of samples and markers relatively inefficient. To address this issue the original implementation of FarmCPU suggested, but did not require, the use of the `big.matrix` data structure from the `bigmemory` package [10] as an alternative to the base `R data.frame` for the storage and manipulation of the marker matrix.

`big.matrix` is a C++ data structure that returns a pointer to the R process. The structure and its pointer can also be stored in binary files and shared across processes to facilitate reuse and parallel computing. While `big.matrix` allows the user to exceed R’s internal limit on object size, *sufficient RAM to store the dataset must still be available* [10]. Thus, `bigmemory` is a useful tool for enabling the analysis of large datasets but is still subject to the physical limitations of the user’s hardware.

The original implementation of FarmCPU has two shortcomings with respect to its handling of a `big.matrix` due to the details of the C++ implementation of the `big.matrix`.

First, a `big.matrix` can store only one of four (`char` [1 byte], `short` [2 bytes], `int` [4 bytes], `double` [8 bytes]) data types at once. Marker scores for GWAS are typically represented as 0, 1, and 2, counting the number of minor alleles that an individual possesses at a single locus. Missing marker scores are often imputed as the mean non-missing marker score (or twice the minor allele frequency). Because minor allele frequency is in the interval (0, 0.5], imputed marker scores will be in the interval (0,1.0]. Therefore, marker scores are a mix of integers and real numbers. Base R seamlessly converts integers into real numbers and, in a dataframe, can store both types. However, real numbers improperly stored in a `char, short`, or `int` type `big.matrix` will be truncated, causing an inflated number of homozygous major individuals. The manual for FarmCPU recommends the use of a `char` type `big.matrix;` this is only appropriate in the absence of missing data—a condition that is not met in real data without imputation. Because storage of a double requires 8 times the space of a `char`, in the presence of missing missing marker data, FarmCPU requires 8× more memory than reported.

Second, FarmCPU takes subsets of the `big.matrix` to handle samples with missing phenotypic data. There are two methods for subsetting a `big.matrix` that also return a `big.matrix`, but they have very different consequences for memory usage. The first option uses the `sub.big.matrix()` function to extract a continuous subset (or block) of the `big.matrix`. This can be handled efficiently using pointers, requiring no copying of the original matrix. The second option uses the `deepcopy()` function to extract a discontinuous subset, creating a new, potentially smaller `big.matrix`. In the worst case that the entire `big.matrix` is copied, `deepcopy()` doubles the memory used to store the data if the original `big.matrix` is not discarded. FarmCPU uses `deepcopy()` to remove markers scores for samples with missing phenotypes without releasing the memory used by the original marker matrix.

FarmCPUpp exclusively uses the `big.matrix` as the data structure to represent the marker matrix. This allows the marker matrix to exceed R’s internal limits on object size, rapidly reload a `big.matrix` into RAM, and easily share the `big.matrix` between different processes. Due to the implementation of single marker regression, FarmCPUpp only accepts a `double`-type `big.matrix` and will throw an error if given any other type. This ensures accurate representation of the marker scores. FarmCPUpp also declines to make any copies of the marker matrix using sub.`big.matrix()` or `deepcopy()`. Instead, missing data is handled by removing the appropriate marker scores using compiled code when that marker is tested. While this strategy would be inefficient in R, it is efficient in compiled code and eliminates the need to make expensive copies of the marker matrix either in RAM or on disk.

### Compiled single-marker regression

As mentioned above, FarmCPU uses the computationally efficient GLM, which has an asymptotic running time of *O*(*nm*) [9]. However, because FarmCPU optimizes the set of pseudo-QTNs used as covariates and iterates until this set converges, the actual runtime of FarmCPU will be proportional to the number of iterations which is a function of the dataset. FarmCPU splits the model matrix into two parts: a constant part including covariates and pseudo-QTNs and a variable part containing the marker scores for the current marker. FarmCPU implements this in R, and it is sufficient to perform GWAS on reasonably large datasets in less than one day. However, this does not scale well with sample size or number of markers. FarmCPUpp uses the same strategy but implements it in C++ using the `Rcpp` [11] and `RcppEigen` [12] packages.

### Parallelization

Single-marker regression assumes that tests for each marker are mutually independent. Therefore, another complementary strategy for increasing the speed of single-marker regression is parallel computing. The version of FarmCPU (v1.02) used for benchmarks has code for parallelized single-marker regression and an option to choose the number of cores used for analysis, but use of this code will throw an error when the first single-marker regression loop is reached. FarmCPUpp uses the `RcppParallel` [13] package and the `ncores.glm` parameter to provide parallelized single-marker regression.

FarmCPU’s model selection chooses the optimal number of pseudo-QTNs to add as covariates in each iteration. This optimization is optional but recommended by the authors. This step is not as large of a bottleneck as single-marker regression, but it can take a significant amount of time when the number of models to test, sample size, and/or number of pseudo-QTNs is large. FarmCPUpp uses the foreach [14] package and the ncores.reml parameter to parallelize this step.

### Feature comparison

FarmCPUpp implements only the core GWAS-related functionality of FarmCPU. Some functionality has been simplified, such as the choice of functions to use for single-marker regression because FarmCPUpp has only one option. Other functions, such as power and FDR tests, are not implemented by FarmCPUpp. Full information on each parameter can be found in the documentation.

FarmCPUpp also implements functions to generate manhattan and QQ plots. Unlike FarmCPU, these are separate from the GWAS function. This allows the user to control base R plotting variables. Sensible default values have been selected for each function.

### Simulations

Markers were simulated in PLINK v1.9 [15] [16] using the `--simulate-qt` flag. All marker sets were simulated with reference allele frequencies in [0.05, 0.5], *D*′ = 0.75, and dominance deviations of 0. The per-QTL additive variance was 1/*m*, where *m* is the number of markers, according to PLINK’s internal upper bound.

Two series of datasets were simulated. The first series had 60,000 markers and sample sizes of 500, 1,000, 2,500, 5,000, 7,500, or 10,000. The second series had 10,000, 50,000, 100,000, 500,000, 1,000,000, or 5,000,000 markers and a sample size of 1,000.

Following [17], phenotypes with a heritability of 0.3 were simulated for each dataset using 3,000 QTNs with effect sizes of 0.9^*k*^, where *k* is the *k*th QTN. Phenotypes were constructed by summing up the QTN effects for each individual and adding i.i.d. normal deviates with mean 0 and variance 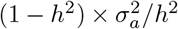, where *h*^2^ is the heritability and 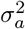 is the variance of the genetic effects.

### Benchmarks

All benchmarks were run on a Ubuntu 14.04 server with 44 Intel Xeon E5-2697 v3 @ 2.60 GHz processors, 377 GB of RAM, and a 4.6 TB HDD. All timing runs were performed with the optimum option for pseudo-QTN selection, the default options for number of pseudo-QTNs, a maximum of 20 iterations, and 1 core unless otherwise stated. The size of bins varied with the number of markers according to Table 1.

**Table 1:** Bin sizes used for model selection at different marker densities.

| Simulation | Bins |
| --- | --- |
| 10,000 markers | 500, 1,000, 1,500, 2,000 |
| 50,000 markers | 2,500, 5,000, 7,500, 10,000 |
| 60,000 markers | 2,500, 5,000, 7,500, 10,000 |
| 100,000 markers | 5,000, 10,000, 15,000, 20,000 |
| 500,000 markers | 25,000, 50,000, 75,000, 100,000 |
| 1,000,000 markers | 50,000, 100,000, 150,000, 200,000 |
| 5,000,000 markers | 250,000, 500,000, 750,000, 1,000,000 |

## Results

### Correctness

The final results from all 12 simulated datasets were compared using the `all.equal()` function in R. FarmCPU reports p-values and effect estimates for each SNP, so we compared only these two measures even though FarmCPUpp also reports the standard errors and t-values for each SNP. FarmCPU and FarmCPUpp produced identical results for effect estimates and p-values.

### Benchmarks

The performance of FarmCPU and FarmCPUpp was tested first on simulated datasets of 60,000 markers with 3,0 markers chosen as QTNs and a heritability of 0.3. Sample size varied from 500 to 10,000. Both packages scale well with increasing sample size (Figure 1a); at a sample size of 10,000, FarmCPUpp is faster by 159 s. In each iteration the FarmCPU algorithm adds pseudo-QTNs to the model as covariates for single-marker regression. Increased model size has a stronger effect on FarmCPU than on FarmCPUpp (Figure 1b). At a sample size of 10,000, the longest single-marker regression time (most complex model) for FarmCPUpp is as fast as the shortest single-marker regression time (least complex model) for FarmCPU.

**Figure 1:**
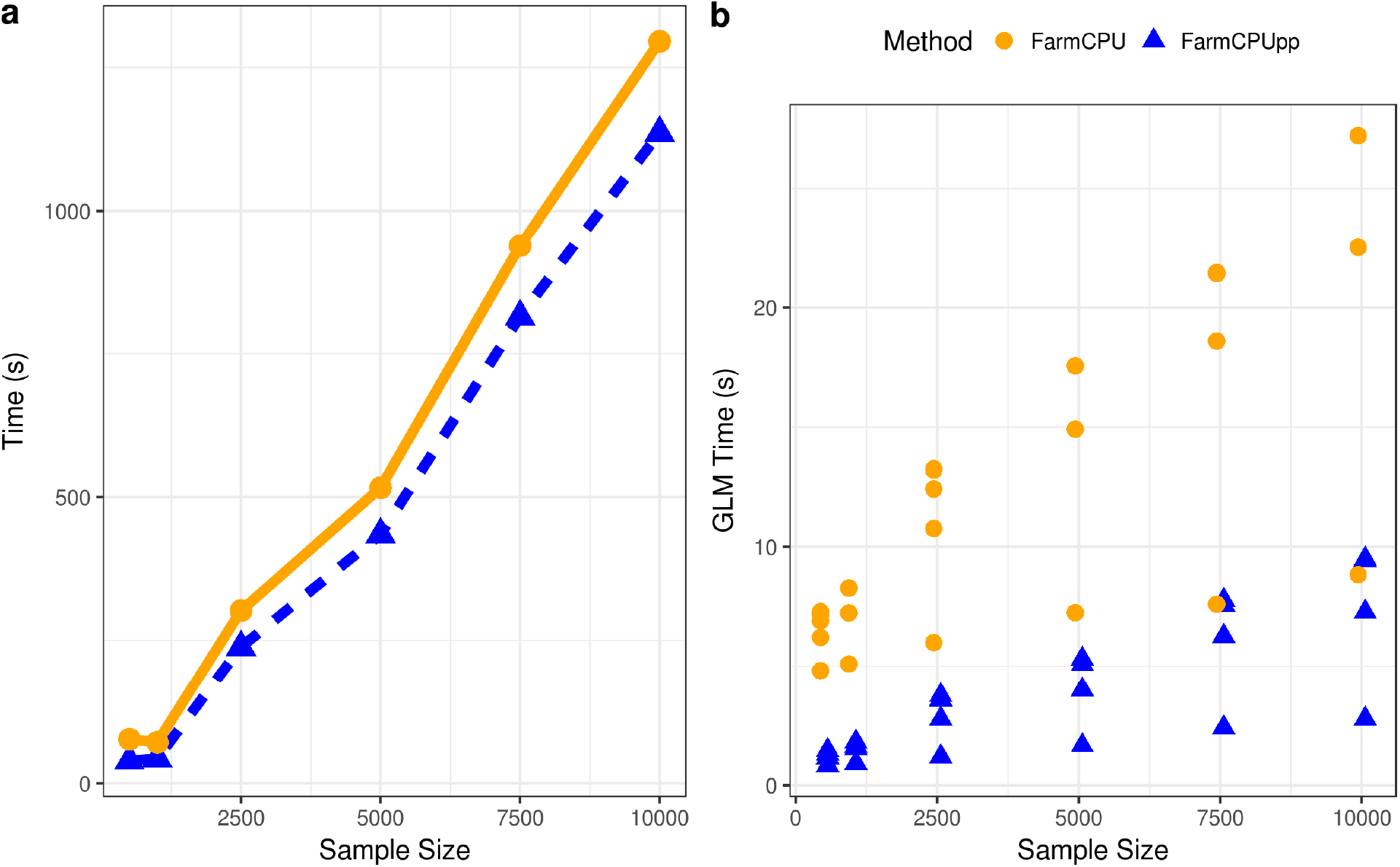
Effects of sample size on the runtime of FarmCPU and FarmCPUpp. **a**. Total runtime in seconds. **b**. Time for single-marker regression in seconds. Dots represent the time spent in each iteration at the indicated sample size.

By contrast, the number of markers has a much stronger effect on the runtimes of both packages. These datasets were simulated with a sample size of 1,000 and numbers of markers that varied from 10,000 to 5,000,000. Phenotypes were simulated with 3,000 QTN and heritabilities of 0.3. With small samples sizes, the performance of both packages is similar at low numbers of markers (Figure 2a). However, FarmCPU scales much more poorly as the number of markers increases. With 5,000,000 markers, FarmCPU takes almost 2 hours to complete while FarmCPUpp takes only 30 minutes. This reduction in runtime is largely attributable to reductions in the time spent on single-marker regression (Figure 2b). With 5,000,000 markers, FarmCPU performs single-marker regression in 517-832 seconds while FarmCPUpp performs the same analysis in 137-185 seconds, a 74-78% reduction in runtime.

**Figure 2:**
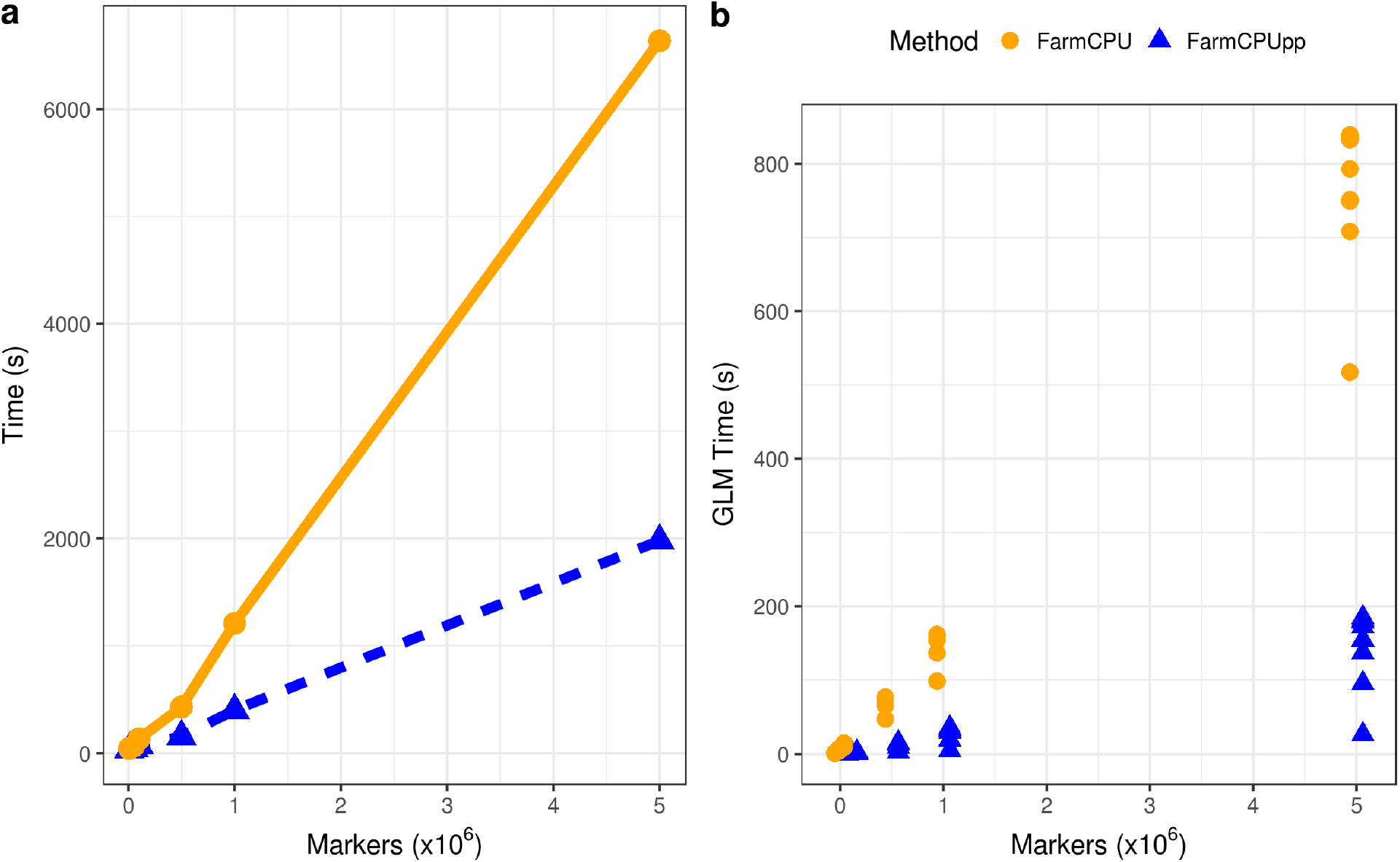
Effects of number of markers on the runtime of FarmCPU and FarmCPUpp. **a**. Total runtime in seconds. **b**. Time for single-marker regression in seconds. Dots represent the time spent in each iteration at the indicated number of markers.

Finally, we tested the performance of both packages on a real dataset described in [18] that consisted of measurements for days to silk on 4,890 recombinant inbred lines (RILs) in the maize nested association mapping (NAM) population [19] (Figure 3). The first three principal components of the marker matrix were used to control for population structure, and 2,452,207 markers were tested. We also used different numbers of cores in FarmCPUpp. As already shown in Figure 1, FarmCPUpp scales slightly better with increasing sample sizes, and using the full dataset this difference is ~6.5 hours or 2.5 × the runtime of FarmCPUpp. This difference represents the advantage of using C++ rather than R for single-marker regression. The use of multiple cores (Figure 3) further decreases the runtime of FarmCPUpp to a minimum observed runtime of ~51 minutes with 16 cores.

**Figure 3:**
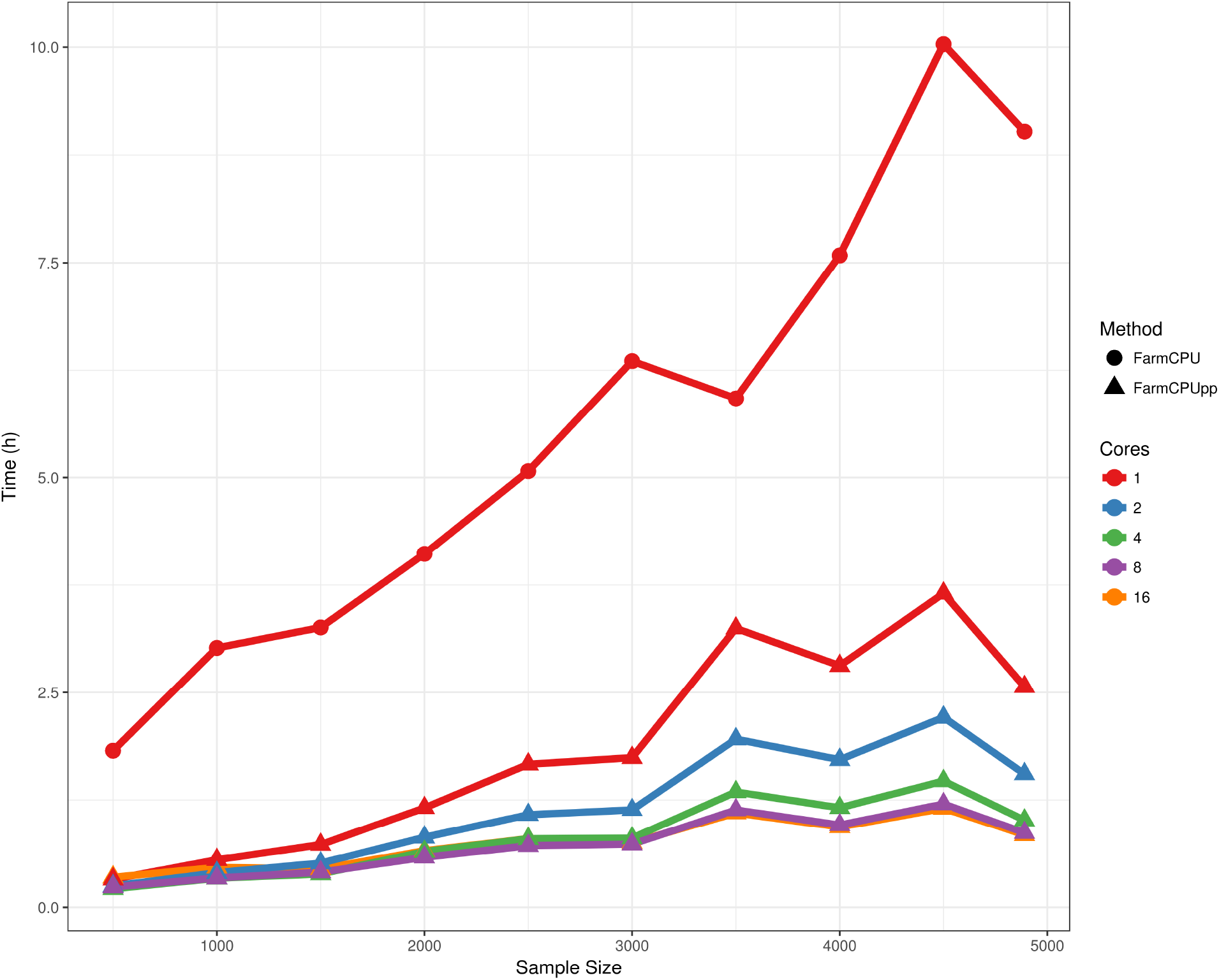
Effects of sample size and number of CPU cores on the runtime of FarmCPU and FarmCPUpp in seconds. The full dataset has 4,890 samples, 2,452,207 markers, and 3 population structure covariates.

## Discussion

GWAS datasets are rapidly growing due to improved phenotyping and genotyping methods, and it is now common to perform GWAS on datasets consisting of thousands of samples and millions of markers. Even though the GLM used in FarmCPU is linear with respect to number of samples and markers, runtimes rapidly increase when both dimensions of the data are increasing. This greater runtime may be acceptable when performing a single GWAS, but GWAS on multiple phenotypes are also common and long runtimes may be prohibitive in these situations. This growth poses a computational challenge to existing methods and software. We have presented FarmCPUpp, an improved version of the FarmCPU method, that uses compiled single-marker regression code and parallel computing to reduce the memory requirements and runtime, especially for large datasets.

The performance of FarmCPU and FarmCPUpp is comparable for datasets up to 10,000 samples by 60,000 markers or 1,000 samples by 1,000,000 markers. Runtimes may still be quite different below these thresholds depending on the dimensions of the data. FarmCPUpp achieves this speedup without parallel processing, although further speedups can be achieved if multiple cores are available. Because the interface of FarmCPUpp is nearly identical to FarmCPU’s, it is not difficult to integrate FarmCPUpp into an existing analysis.

In summary, FarmCPUpp makes improvements on the original FarmCPU R code through the use of C++ and parallel computing. This decreases the memory requirements and runtime while maintaining the R interface of the original software. FarmCPUpp is available as an R package on GitHub at https://github.com/amkusmec/FarmCPUpp.

## Acknowledgments

This material is based upon work supported by the National Science Foundation (grant number 1027527) to P.S.S.

## Author Contributions

A.K. designed and wrote the modified software. A.K. and P.S.S. wrote the manuscript.

